# Characterization of the gene repertoire and environmentally driven expression patterns in Tanner crab (*Chionoecetes bairdi*)

**DOI:** 10.1101/2021.05.10.443482

**Authors:** Grace Crandall, Pamela C. Jensen, Samuel J. White, Steven Roberts

**Author notes:** Corresponding author: Steven Roberts.

## Abstract

Tanner crab (*Chionoecetes bairdi*) are an economically important species that is threatened by ocean warming and Bitter Crab Disease, which is caused by an endoparasitic dinoflagellate, *Hematodinium*. Little is known about disease transmission or its link to host mortality, or how ocean warming will affect pathogenicity or host susceptibility. To provide a transcriptomic resource for the Tanner crab, we generated a suite of RNA-seq libraries encompassing pooled hemolymph samples from crab displaying differing infection statuses and maintained at different temperatures (ambient (7.5°C), elevated (10°C), or decreased (4°C)). After assembling a transcriptome and performing a multifactor differential gene expression analysis, we found genes influenced by temperature in relation to infection, and detected some of those genes over time at the individual level using RNAseq data from one crab. Biological processes associated with those genes include lipid storage, transcription, response to oxidative stress, cell adhesion, and morphogenesis. Alteration in lipid storage and transcription provide insight into how temperature impacts energy allocation in *Hematodinium* infected crabs. Alteration in expression patterns in genes associated with morphogenesis could suggest hemocytes were changing morphology and/or type in response to temperature. This project provides insight into how *Hematodinium* infection could influence crab physiology as oceans warm.

## Introduction

Southern Tanner crab (*Chionoecetes bairdi*), hereafter referred to as Tanner crab, is an important species for commercial, recreational, and subsistence fishing. They occur in relatively shallow waters along the continental shelf, from the Bering Sea in Alaska through coastal Oregon (Jadamec et al. 1999), and are found at temperatures ranging from −2°C to 9°C (Nielsen et al. 2007). There are many threats to Tanner crab during all phases of their life, among them ocean warming and disease. One disease, bitter crab disease (BCD), has been termed the “principal threat” to Alaska Tanner crab stocks by the Alaska Department of Fish and Game (Alaska Department of Fish and Game). BCD is caused by a parasitic dinoflagellate of the genus *Hematodinium*, and Tanner crab in southeast Alaska experience high infection rates, approaching 100% in some areas (Meyers et al. 1987, 1990); (Eaton et al. 1991); (Love et al.1993); (Bednarski et al. 2011). *Hematodinium* spp. infects 40+ decapod crustacean species worldwide (Morado 2011), and its spread around the world has followed warming trends in the Atlantic and Pacific Oceans (Morado et al. 2011). This has implications for *Hematodinium* spp. to continue to spread to naive Tanner crab and other crustacean populations with increasing temperature. There are two described *Hematodinium* species (Chatton and Poisson 1931); (Hudson and Shields 1994), and an additional recognized but undescribed species that infects Tanner crab (Jensen et al. 2010).

Crustacean innate immunity centers around the activity of their hemocytes, of which there are three types: hyaline cells (phagocytosis), granular cells (melanization, antimicrobial peptides, cytotoxicity), and semi-granular cells (encapsulation, early non-self recognition, coagulation) (Lin and Söderhäll 2011). Granular and semi-granular cells are critical in initiating and sustaining the cascade of responses that are involved in the major innate immune responses of the prophenoloxidase (proPO) system and melanization (Cerenius and Soderhall 2004). The immune response begins when host protein recognition proteins (PRPs) recognize non-self pathogen-associated molecular patterns (PAMPs) (e.g., bacterial peptidoglycans), which initiates an immune response that leads to the production of melanin, which binds to nucleophiles on pathogen surfaces and encapsulates pathogens (Cerenius et al. 2008); (Nappi and Ottaviani 2000). There are additional innate immune response pathways, with some of the important processes described in Verbruggen et al. (Verbruggen et al. 2015).

The mode of transmission is unknown, as well as whether *Hematodinium* always causes direct mortality, or if it can be a chronic condition (Gornik et al. 2010) and the extent to which death comes via secondary factors such as the increased risk of predation to lethargic crab (Butler et al. 2014). The disease alone is concerning, but with the mounting threat of increasing ocean temperatures, it is important to understand how these two threats will interact to affect Alaska’s Tanner crab populations. In the Atlantic Ocean in the closely related species *Chionoecetes opilio*, infection with *Hematodinium* has been correlated with increased temperatures (Shields et al. 2007). In other species and marine ecosystems, it has been documented that increasing temperature decreases the hosts’ ability to mount an immune defense against pathogens (Bruno et al. 2007), increases the prevalence of disease (Groner et al. 2018), and can increase a pathogen’s ability to grow and reproduce, thus enhancing infectivity potential (Harvell et al. 2002).

Although *Hematodinium* spp. transmission remains a black box, research has uncovered some potential pathways. It is possible that moribund infected hosts shedding infective-stage *Hematodinium* dinospores in the water column could infect other crab. The dinospores may enter through tears in the exoskeleton during molting, or through another method unassociated with molting, though it is unknown how the dinospores could penetrate the exoskeleton (Rowley et al. 2015); (Meyers et al. 1990). It is not well understood whether there is a host immune response to the *Hematodinium* spp., as there is no evidence of parasite encapsulation, and it is thought that *Hematodinium* spp. may be able to conceal itself from the host immune system through molecular mimicry or evasion (Rowley et al. 2015).

In crustacean hosts with BCD, there are some outward signs that aid with diagnosis, as well as technical methods of detection. BCD causes milky, opaque hemolymph, pinkish discoloration and cooked-like appearance to the carapace, and lethargic behavior (Eaton et al.1991). A healthy crab has clear hemolymph, but in infected crabs, *Hematodinium* proliferates throughout the hemal spaces, causing the hemolymph to appear milky. Eventually, all organ systems become impacted and ultimately, the parasite outcompetes the host for nutrients and oxygen, leading to host lethargy and subsequent suffocation. While BCD is not harmful to humans, it renders their meat chalky and bitter, which makes them unmarketable. As such, the Tanner crab fishery has suffered direct economic loss (Meyers et al. 1987, 1990). Other infections can present similar gross signs, such as the milky hemolymph syndrome (MHS) described in *C. opilio*, which causes opaque hemolymph in response to infection with a bacilliform virus (Kon et al. 2011), so additional detection methods should be used for confirmation of *Hematodinium* spp. presence. Common methods are conventional PCR (Jensen et al. 2010) or quantitative PCR (Crosson 2011), to detect *Hematodinium* spp. DNA, and examining hemolymph smears or other host tissues to directly detect *Hematodinium* spp. *in situ* (Shields 2017).

We aimed to investigate how temperature and infection with *Hematodinium* spp. impact crab physiology. We generated a *Chionoecetes bairdi* transcriptome and used it for differential gene expression analysis in infected and uninfected crab held at three temperatures and sampled at three time points. For exploratory purposes, we also were able to visualize gene expression patterns at the individual level over time using an individual crab RNAseq data. Our results demonstrate how environmental conditions impact crab gene expression and provide insight as to how *Hematodinium* sp.-infected southeast Alaska Tanner crab may fare as ocean temperatures continue to rise.

## Methods

### Tanner crab collection and disease status determination

In late October 2017, 400 morphometrically immature male Tanner crab were collected using crab pots by the Alaska Department of Fish and Game in Stephen’s Passage, near Juneau, Southeast Alaska. Immature male crab were chosen to avoid having too many additional variables in the experiment, and had a carapace width to claw height ratio < 0.18 mm (Tamone et al. 2007). Stephen’s Passage was selected because its long term *Hematodinium* sp. rate of infection in Tanner crab is ~50% (ADF&G unpub. data). The crab were transported to Ted Stevens Marine Research Institute (TSMRI, NOAA facility, Juneau, AK) and placed in flow-through seawater tanks at 7.5°C, the benthic water temperature in Stephen’s Passage at the time of capture.

From each crab, 200 ul of hemolymph was withdrawn and preserved in 800 μl 95% ethanol for PCR detection of *Hematodinium* sp. infection. For extraction of total genomic DNA, 200 μl of ethanol-preserved hemolymph was centrifuged to pellet the solids, the supernatant discarded, and the pelleted material air-dried and processed as tissue. DNA was extracted following Ivanova et al. (2006) using invertebrate lysis buffer and modified by performing 2 washes with Wash Buffer, and adjusting eluted DNA (50 μl) to 10 mM Tris-Cl, pH 8.0, and 0.1 mM EDTA. Genomic DNA was subjected to 2 rounds of PCR with 2 different primer pairs designed for the small subunit (SSU) rRNA gene of *Hematodinium* spp.: Univ-F-15 and Hemat-R-1654 (Gruebl et al. 2002) and Hemat 18Sf and Hemat 18Sr (Bower et al. 2004). Reaction aliquots were pooled post-PCR and visualized on ethidium bromide-stained 2% agarose gels. Samples were scored as positive when both *Hematodinium* sp. bands of the expected size were visible on the gel, negative when neither fragment amplified.

### Experimental design

The crab were allowed to acclimate at 7.5°C for 9 days. At the end of the acclimation period, of the crab that appeared to have recovered from capture stress, 180 were selected for temperature treatments, such that half were infected and half were uninfected as determined by PCR. Twenty crab (10 infected and 10 uninfected) were placed in each of 9 replicate tanks at 7.5°C. Over the course of day 0 through day 2, the temperature in 6 tanks was gradually adjusted to the final experiment temperatures of 10°C (elevated) in 3 tanks and 4°C (decreased) in 3 tanks; the remaining 3 tanks were kept at 7.5°C. Prior to the initiation of the temperature adjustments (day 0), 0.2 ml hemolymph was sampled and preserved in 1200 μl RNAlater (Qiagen) for transcriptomic analyses. Hemolymph was sampled and preserved in RNAlater again after 2 days and at the termination of the temperature trial (day 17).

### RNA Sequencing for Transcriptome Assembly

Hemolymph samples (n=112) were centrifuged for 10 minutes at 14000 g with RNA extracted using Quick DNA/RNA Microprep Plus Kit (Zymo Research) according to the manufacturer’s protocol. Samples (2 μl) were run on Qubit 3.0 using the Qubit RNA HS Kit (Invitrogen) to determine RNA quantification. Samples were pooled to increase complexity of transcriptome as well as in response to limited RNA yield from hemolymph samples. Specifically, a total of 14 libraries were constructed (Supplemental Table S1). RNA was submitted to Northwest Genomics Center at Foege Hall at the University of Washington for library construction and sequencing. Samples were sequenced on an Illumina NovaSeq with 100bp paired-end reads.

### RNA Sequencing for Differential Gene Expression

Four of the 11 eleven libraries above were used for differential gene expression analysis. For differential gene expression all libraries were from sampling day 2 and were pools of 10 individuals (Supplemental Table S1). Two libraries were generated from crabs held at decreased temperature (infected and uninfected) and two libraries were generated from crab held at elevated temperature (infected and uninfected). RNA was submitted to Northwest Genomics Center at Foege Hall at the University of Washington, where RNA-seq libraries (100bp PE) were constructed and sequenced on a NovaSeq 6000 (Illumina). Additionally, for exploratory purposes, RNA was also extracted from a single individual held in ambient conditions and determined to be infected with *Hematodinium* sampled over the three time points (day 0, day 2, and day 17). These three samples were sent to Genewiz, Inc. where RNA-seq libraries were constructed and sequenced (Illumina HiSeq4000; 150bp paired-end) in order to look at the gene expression at the individual level over time.

### Transcriptome Assembly and Annotation

Raw sequence data from the 11 pooled libraries were assessed using FastQC (v0.11.8; (Andrews 2010)) and MultiQC (v1.6; (Ewels et al. 2016)) pre- and post-trimming. Data were quality trimmed using fastp (v.0.20.0; (Chen et al. 2018)) with the “--detect_adapter_for_pe” setting. A transcriptome was *de novo* assembled using Trinity (v2.9.0; (Grabherr et al. 2011); (Haas et al. 2013)). All raw sequencing data is available in the NCBI Sequence Read Archive (SRR11548643 - SRR11548677).

The initial transcriptome assembly was further refined by excluding all sequences within and below the superphylum Alveolata (dinoflagellates). Sequences were functionally annotated and taxonomically categorized with a combination of DIAMOND BLASTx (0.9.26; (Huson et al.2016)) and MEGAN6 (6.18.3; (Huson et al. 2016)). Annotation with DIAMOND BLASTx was run against NCBI nr database (downloaded 20190925). The resulting DAA files were converted to RMA6 files for importing into MEGAN6 with the daa2rma utility, using the following MEGAN6 mapping files: prot_acc2tax-Jul2019X1.abin, acc2interpro-Jul2019X.abin, acc2eggnog-Jul2019X.abin. All eukaryotic sequences, sans those categorized within and below Alveolata, were extracted using MEGAN6 to produce the transcriptome assembly used for all analysis.

Transcriptome “completeness” was assessed with BUSCO (v3.1.0; (Simão et al. 2015;Waterhouse et al. 2018)) using the transcriptome option, metazoa_odb9 database, and AUGUSTUS (v3.3.2; (Stanke and Waack 2003; Stanke et al. 2006a, b, 2008)) with “fly” set as species. TransDecoder (v5.5.0; (Grabherr et al. 2011; Haas et al. 2013)) was used to identify putative open reading frames (ORFs), using BLASTp (v2.8.1+; (Altschul et al. 1990; Camacho et al. 2009)), and HMMER (v.3.2.1; http://hmmer.org/). Trinotate (v3.3.1;(Bryant et al. 2017)) was used to assign functional annotations (gene ontology) to genes using BLASTx (v2.8.1+; (Altschul et al. 1990; Camacho et al. 2009)), RNAMMER (v1.2; http://www.cbs.dtu.dk/services/RNAmmer/), SignalP (v4.1; (Petersen et al. 2011)), tmhmm (v2.0c; (Sonnhammer et al. 1998)) and the longest ORFs identified by Transdecoder.

### Single Nucleotide Polymorphism Identification

Quality-trimmed reads from the four libraries used for differential gene expression (380822, 380823, 380824, 380825) were aligned to the transcriptome assembly with HISAT2 (v2.1.0; (Kim et al. 2019). Variant identification and calls were generated using bcftools (v1.13) ‘mpileup’ and ‘call’ commands (Li 2011; Danecek et al. 2021). Filtering for quality (minimum 30) and raw read alignment depth (minimum 10) was performed using bcftools (v1.13) ‘filter’ command (Danecek et al. 2021). Basic statistics were gathered for each library using bcftools (v1.13) ‘stats’ command (Danecek et al. 2021).

### Differential Gene Expression Analysis

Gene expression differences were assessed by the comparison of RNA-seq data. Specifically, this included four of the libraries used to develop the transcriptome. Kallisto was used to obtain count data for each library and an abundance matrix was then produced using a perl script (abundance_estimates_to_matrix.pl) provided as part of Trinity (v2.8.6) (Grabherr et al. 2011); (Haas et al. 2013). Differential expression of contigs was calculated using a negative binomial GLM in the R package DESeq2 (Love et al. 2014). The read counts were first normalized using the size factors method and fit to a negative binomial distribution. Significantly differential contig expression (Benjamini-Hochberg adjusted p<0.05) between infected and uninfected crabs was determined using the Wald test for significance of GLM terms.

Analyses were performed to address how temperature affects the contigs differentially expressed between infected and uninfected crabs. Differential expression of contigs was calculated between two libraries from uninfected crabs and the two libraries from infected crabs to capture differences due to temperature treatment (decreased or elevated) in a multifactor design formula. From the results of the multifactor differential expression of contigs, a contrast was performed between the two temperature treatments to extract the differentially expressed contigs related to infection with *Hematodinium* that are influenced by temperature. All associated code is available in the corresponding repository.

### Time series gene expression

For exploratory purposes, expression patterns from an infected individual crab held at ambient temperature and sampled at the three time points were examined. Kallisto (Bray et al.2016) was used to obtain count data for each library for the individual crab over three time points and an abundance matrix was then produced using a perl script (abundance_estimates_to_matrix.pl) provided as part of Trinity (v2.8.6) (Grabherr et al. 2011); (Haas et al. 2013). Highly expressed contigs (> 63 counts across the three sampling time points) were used to visualize expression over time (**Figure 3**). The corresponding cladogram was used to determine the clusters of genes that change over time in a similar pattern. Gene enrichment analysis was performed as described below to functionally annotate processes associated with each cluster. Additionally, genes identified as part of the differential expression analysis were identified in these libraries to explore expression over time. All associated code is available in the corresponding repository.

### Enrichment Analysis

Gene enrichment analysis was performed using DAVID v. 6.8 (Huang et al. 2009a); (Huang et al. 2009b). The Uniprot Accession IDs from the annotated crab transcriptome were set as the background. The gene lists were the Uniprot Accession IDs from the differentially expressed contig lists related to infection status that were influenced by temperature, and separately the Uniprot Accession IDs from the contigs identified in the times series analysis.

## Results

### Survival

On day 4 of the temperature trial, a mortality event began in the elevated temperature tanks. By day 10, 95% of the crabs at the elevated temperature had perished. Over the course of the experiment, one crab died in the decreased temperature treatment and three mortalities occurred in the ambient temperature treatment.

### Transcriptome assembly and annotation

Assembly of 143,543,003 base pairs (bp) from the 11 libraries resulted in 78,649 contigs (Supplemental File S2). The median contig length was 1522bp, with an average contig length of 1825 bp and an N50 of 2580 bp (**Table 2**). The resulting assembly had the following BUSCO scores: C:96.5%[S:40.3%,D:56.2%],F:2.2%,M:1.3%,n:978. Of the 78,649 contigs, 48,551 were able to be annotated using the Swiss-Prot database, with 47,731 having corresponding Gene Ontology information (Supplemental Table S3). Of the annotated contigs, 6,191 contigs associated with the Biological Processes GOslim term “stress response” were identified and annotated (Supplemental Table S4). GOslim terms were acquired for the transcriptome and visualized (**Figure 1**).

**Table 2.** Comparison of *de novo* transcriptome assembly statistics between the Tanner crab (this study), white leg shrimp (*Litopenaeus vannamei*) (Ghaffari et al. 2014), and European shore crab (*Carcinus maenas*) (Verbruggen et al. 2015) transcriptomes.

|  | *C. bairdi* (this study) | *L. vannamei* (Ghaffari et al. 2014) | *C. maenas* (Verbruggen et al. 2015) |
| --- | --- | --- | --- |
| Contigs | 78,649 | 87,307 | 212,427 |
| Median contig length | 1522 bp | 429 bp | 380 bp |
| Average contig length | 1825 bp | 1137 bp | 992 bp |
| N50 | 2,580 bp | 2,701 bp | 2,102 bp |
| BUSCO | C:96.5%[S:40.3%,D:56.2%], F:2.2%,M:1.3%,n:978 | C:98.1%[S:72.7%,D:25.4%],F:0.9%,M:1.0%,n:978 | C:95.7%[S:57.0%,D:38.7%], F:3.6%,M:0.7%,n:978 |

**Figure 1.**
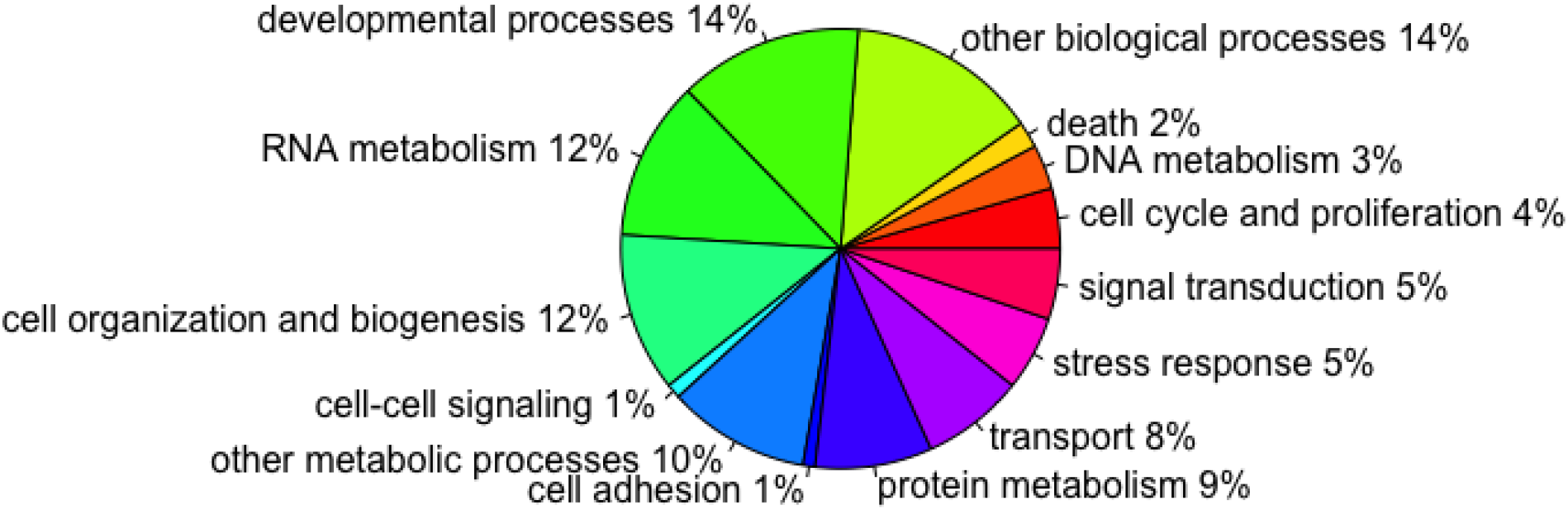
The proportion of Biological Processes GOslim terms for the biological processes identified in the crab transcriptome. Contigs have multiple GO terms, and as such fall under multiple GOslim terms.

### Single Nucleotide Polymorphisms

A total of 79,753 SNPs were cumulatively identified across all libraries (380822, 380823, 380824, 380825). At least one SNP was identified in 17,680 transcripts (22.74% of transcriptome assembly), with a maximum of 384 SNPs identified in a single transcript, and a mean of 4.51 SNPs in any transcript having a SNP. Variant calling identified 50,173 transitions and 30,371 transversions combined in all four libraries. Assigning transcripts with SNPs to Biological Process GOslim terms revealed assignments to each of the 72 GOslim categories. Excluding assignments to the generic “biological process” category (55.53% of all GO terms), “photosynthesis” (0.00017% of all GO terms) and “nitrogen cycle metabolic process” (0.0018% of all GO terms) were the least represented GOslims, while “cellular nitrogen compound metabolism” (6.08% of all GO terms) and “biosynthetic process” (5.23% of all GO terms) were the most represented GOslims (Supplemental Figure S9; Supplemental Table S10).

### Differential Gene Expression

A total of 408 differentially expressed contigs were identified, 341 of which were annotated when considering temperature and infection status. Of these 408 contigs, a majority (357) were expressed at an elevated level in infected crabs (Supplemental Table S5). A total of 123 of the 408 differentially expressed contigs related to infection with *Hematodinium* sp. were influenced by temperature treatment, 103 of which were annotated. There were 70 contigs expressed at an elevated level and 53 expressed at a decreased level in the elevated temperature treatment (**Figure 2**; Supplemental Table S6). There were 4 primary expression clusters of differentially expressed contigs. While there were no significantly enriched processes as determined by DAVID, prominent biological processes of contigs expressed at a relative higher level at elevated temperature (Clusters 1 and 3) include cell adhesion, activation of innate immune response, morphogenesis, cell differentiation, and metabolism. Prominent biological processes of contigs expressed at a relative lower level at elevated temperature (Clusters 2 and 4) include lipid storage, development, response to oxidative stress, and metabolism.

**Figure 2.**
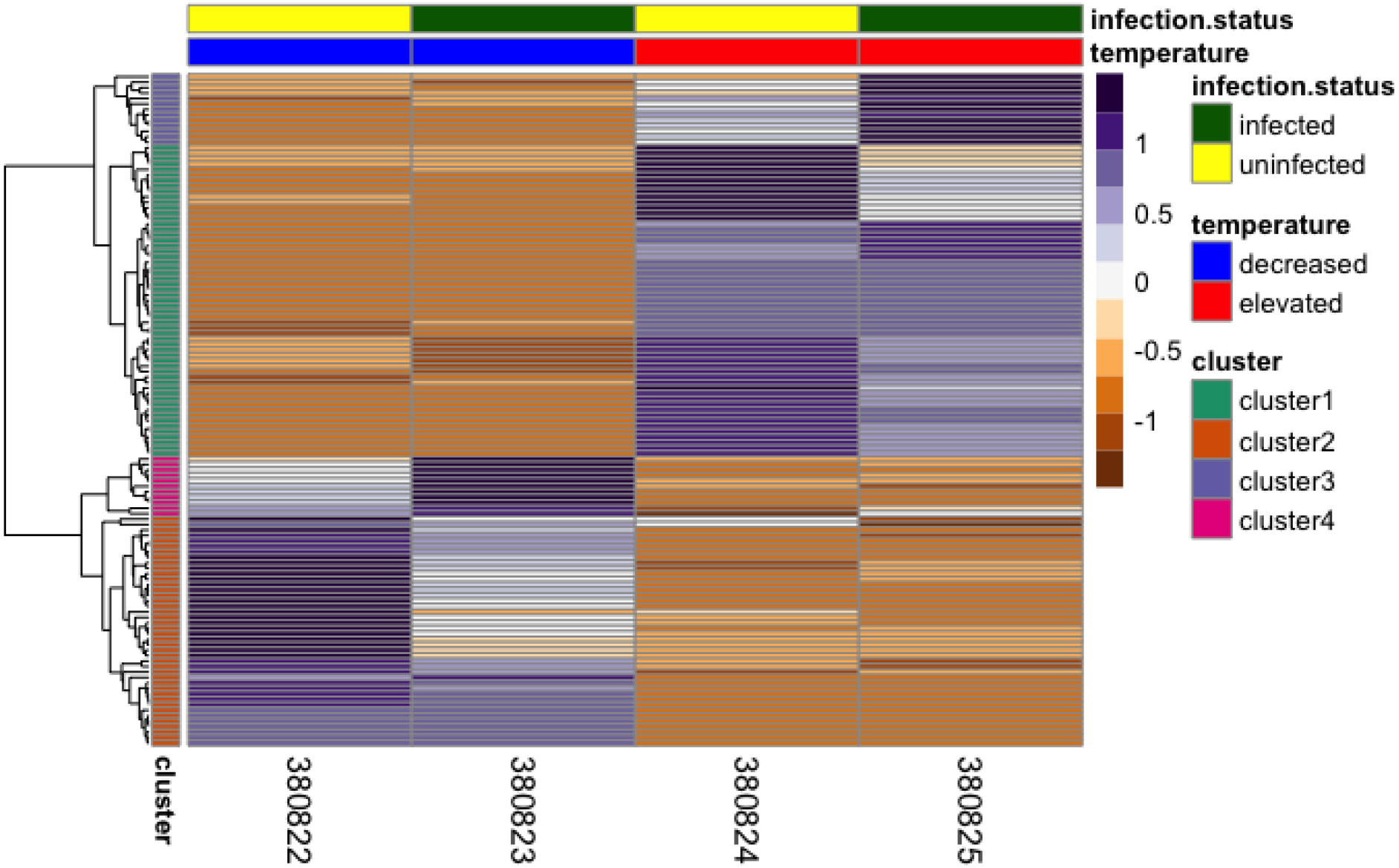
Differentially expressed contigs (n=123) associated with *Hematodinium* sp. infection and driven by temperature treatment. Each row is a contig, with blue/purple coloration indicating higher expression, and orange/brown coloration indicating lower expression. The coloration correlates with the scaling values across the rows (contig expression counts). The scales are Z scores, with above zero indicating higher expression, and below zero indicating lower expression. Library IDs are on the X-axis. The cladogram on the left groups contigs that have similar expression levels. The cluster bar annotates the different groups of contigs based on the cladogram.

### Time series gene expression

A total of 13,954 contigs were characterized in a single crab over the three time points (Supplemental Table S7). There were 6 clusters identified and associated with expression patterns (**Figure 3**). A majority of the contigs ended up with relatively decreased expression by day 17 (Clusters 1,2,3). The contigs in Cluster 2, (and to a lesser degree Clusters 1 and 6) demonstrated relative elevated expression at day 2. While there were no significantly enriched processes as determined by DAVID, biological processes represented in clusters 1-3 are associated with a wide range of functions including development, transcription, and phagocytosis.

**Figure 3.**
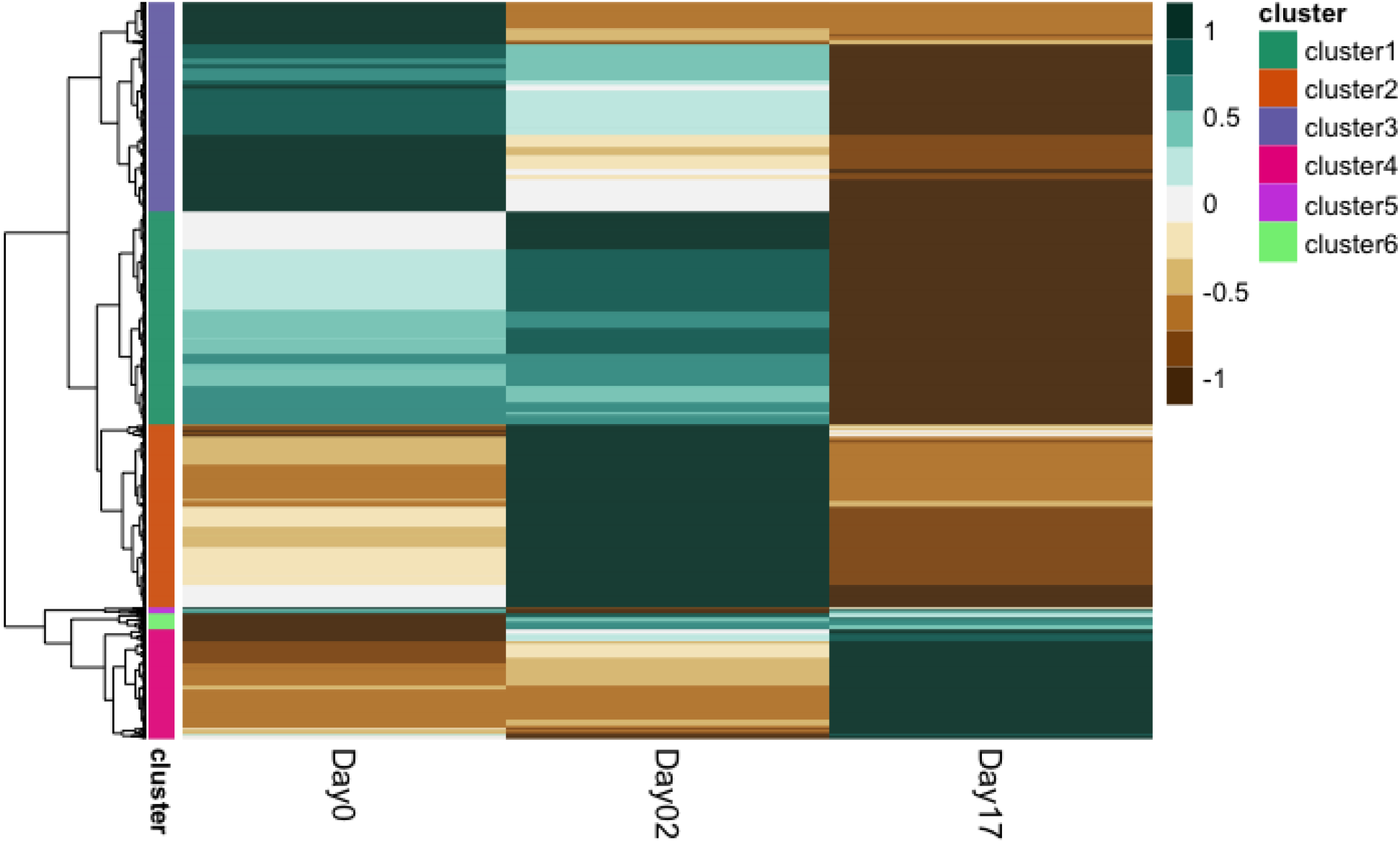
Differentially expressed contigs (n=13,954) in an individual *Hematodinium* sp.-infected crab over time. The individual crab was held at ambient temperature. Each row is a contig, with the blue/green coloration indicating higher expression, and orange/brown coloration indicating lower expression. The coloration correlates with the scaling values across the rows (contig expression counts). The scales are Z scores, with above zero indicating higher expression, and below zero indicating lower expression. The cladogram on the left groups contigs that are more similar. The cluster bar annotates the different groups of contigs based on the cladogram.

### Temperature-influenced differentially expressed contigs in the individual crab

Of the 123 differentially expressed contigs associated with infection status that are influenced by temperature in the pooled libraries, 74 were present in the individual crab over time (Supplemental Table S8). Of the differentially expressed contigs (**Figure 3**), 12 were present in cluster 1 and 30 of the differentially expressed contigs present in cluster 2. There were 29 differentially expressed contigs present in cluster 3, 2 present in cluster 4, and 1 present in cluster 5.

### Enrichment Analysis

There were no significantly enriched biological processes in any of the analyses.

## Discussion

In this study, we have made progress towards understanding the mechanisms behind the influence of temperature on infection with the endoparasite *Hematodinium* sp. on the immune response of one of its hosts, *Chionoecetes bairdi*. We produced and annotated a *Chionoecetes bairdi* hemolymph transcriptome from *Hematodinium* sp.-infected and uninfected crab exposed to different temperature treatments. Using the transcriptome and libraries of interest, we were able to identify genes that were differentially expressed between *Hematodinium* sp.-infected crabs and uninfected crabs as well as identifying which of these genes were significantly influenced by temperature. Furthermore, we identified 79,753 single nucleotide polymorphisms (SNPs) within the data providing a valuable genomic resource for future genetic studies in this system.

### Transcriptome Assembly and Annotation

The Tanner crab hemolymph transcriptome is comparable to transcriptomes of the whiteleg shrimp (*Litopenaeus vannamei*) (Ghaffari et al. 2014) and the European shore crab (*Carcinus maenas*) (Verbruggen et al. 2015), with the shrimp transcriptome being more similar in size than that of the shore crab to the *Chionoecetes bairdi* transcriptome. Ghaffari et al. (Ghaffari et al. 2014) assembled the shrimp transcriptome from RNA-seq data from the abdominal muscles, hepatopancreas, gills, and pleopods of one male shrimp. Verbruggen et al. (Verbruggen et al. 2015) assembled the European shore crab transcriptome from RNA-seq data from 12 pooled libraries of 12 tissues and organs from adult males and females.

As part of the Tanner crab transcriptome, 6,191 contigs associated with stress response were identified and annotated (Supplemental Table S4). Of the expressed genes, there were proPO, phenoloxidase, lozenge, peroxinectin, and serine protease, among others, that are involved in the prophenoloxidase system and melanization pathway. Lozenge regulates the expression of proPO, which is a component of the proPO pathway in crustacean innate immunity (Tang 2009; Verbruggen et al. 2015), while peroxinectin contributes to the adhesion of hemocytes to pathogens (Liu et al. 2005); (Verbruggen et al. 2015). There were also genes present that contribute to other innate immune responses, such as the toll-like receptors, which bind to PAMPs (Kingsolver et al. 2013); (Verbruggen et al. 2015). Additionally, dicer-1 was identified, which is part of the RNAi pathway involved in antiviral innate immune response (Wang et al. 2014); (Verbruggen et al. 2015). And as part of the endocytosis pathway, Rab GTPases and others were detected, which aid in phagocytosis and macropinocytosis, and the elimination of pathogens (Egami 2016).

### Differential Gene Expression

As might be expected when looking at differentially expressed genes in infected crab hemolymph, a majority (87%) were elevated, likely indicative of a stimulated immune response. Among genes that were differentially expressed between *Hematodinium* sp.-infected crabs and uninfected crabs there were unc-112-related protein (Fit1), signal peptide peptidase-like 2b (SPPL2B), and eukaryotic translation initiation factor 2-alpha kinase 4 (EIF2AK4) expressed, which may each have roles in immune function. In *L. vannamei*, unc-112-related protein was detected in the immune transcriptome, and this protein has the role of cell adhesion (Robalino et al. 2007). Signal peptide peptidase-like 2b was detected, and while it has several potential functions, it may play a role in the regulation of innate immune response, as described in the annotated list of differentially expressed contigs in the supplemental material. The eukaryotic initiation factor 2-alpha contributes to the immune response of *L. vannamei* to the white spot syndrome virus (Xu et al. 2014). While they have not been studied in Tanner crab, these processes are likely conserved across crustacean species.

When we looked further into genes that were differentially expressed and only considered those influenced by temperature, about half (56%) had elevated expression in infected crab hemolymph in the elevated temperature treatment (**Figure 2**; Supplemental Table 6). In the clusters of contigs that were highly expressed in the elevated temperature treatment crabs (clusters 1 and 3), the biological processes of transcription were prominent and associated with Histone deacetylase complex subunit Sin3a, a gene also expressed higher in uninfected crabs. While this particular gene is best described in humans, some of its functions relate to activation of the innate immune response, indicating that expression of this gene is substantially impacted in infected crabs. Generally speaking, the evidence of several genes involved in transcription at elevated temperature is consistent with an overall increase in metabolism that would be accompanied by the increased transcriptional activity. While less expected, an observed alteration in expression patterns in genes associated with morphogenesis could suggest hemocytes were changing morphology and/or type in response to temperature. For contigs that were highly expressed in the crabs held at decreased temperature (clusters 2 and 4), the associated biological processes included metabolism, lipid storage, and response to oxidative stress. Further analysis of/study of alteration in lipid storage and metabolism could provide insight into how temperature impacts energy allocation in *Hematodinium* sp.-infected crabs. The elevated expression of response to oxidative stress in crab challenged by *Hematodinium* sp. infection and in the decreased temperature treatment could indicate that infection with *Hematodinium* sp. may make the crab more prone to other stresses such as oxidative stress, which can result in cell and tissue damage. This could also make the crab weaker in terms of their ability to fight off *Hematodinium* sp.

### Temperature-influenced differentially expressed contigs in the individual crab

One process that we were able to gain more insight into with the time series analysis was the regulation of the JAK-STAT cascade. The JAK-STAT cascade is part of the crustacean immune response and has been demonstrated to activate in response to the White Spot Syndrome virus infection in shrimp (Verbruggen et al. 2015). This process and the pattern of expression over time could indicate that in the initial days of infection with *Hematodinium*, the crab was able to launch an immune response, but over time the crab weakened and was unable to continue the immune response by the end of the experiment on day 17, although protein analysis should be performed to get a more complete picture. A gene with a corresponding expression pattern was RNA polymerase II promoter. The general decrease of expression over time could be indicative of energy allocation changes in an infected crab over time. Specifically, a taxed immune system might negatively impact the overall ability of a crab to maintain general transcription. The decrease over time of expression of contigs related to cell proliferation and morphogenesis, among other processes, could also indicate an overall depression of a crab’s physiological status.

## Conclusions

The high prevalence of BCD in Alaska is an ongoing threat, and with increasing ocean temperatures as a result of climate change, understanding the interaction of temperature and disease and its effect on the crabs is important. Here we have just started to reveal the gene expression response in the Tanner crab which has provided important insight into immune response and energy allocation. Further, the publicly available assembled *Chionoecetes bairdi* transcriptome provides a valuable resource for future research in genomics for Tanner crab and other decapods.

## Declarations

### Availability of Data and Material

The datasets generated during and/or analyzed during the current study are available in the RobertsLab/paper-tanner-crab repository, available at 10.5281/zenodo.4563060. All raw sequencing data is available in the NCBI Sequence Read Archive (SRR11548643 - SRR11548677).

### Code Availability

Code is available at 10.5281/zenodo.4563060.

### Author Contributions

G.C. performed RNA extractions for sequencing, data analysis and interpretation, and manuscript writing and preparation for publication. P.J. contributed to project and experimental design and performed sample collection from live crab, qPCR, and manuscript writing and editing. S.W. assembled and annotated the transcriptome, and provided manuscript writing and editing. S.R. contributed to the experimental and project design, guidance throughout data analysis and interpretation, and manuscript writing and editing.

